# Landscape composition mediates suppression of major pests by natural enemies in conventional cruciferous vegetables

**DOI:** 10.1101/2021.02.04.429816

**Authors:** Jie Zhang, Shijun You, Dongsheng Niu, Karla Giovana Gavilanez Guaman, Ao Wang, Hafiz Sohaib Ahmed Saqib, Weiyi He, Yuan Yu, Guang Yang, Gabor Pozsgai, Minsheng You

**Affiliations:** State Key Laboratory of Ecological Pest Control for Fujian and Taiwan Crops, Institute of Applied Ecology, Fujian Agriculture and Forestry University, Fuzhou 350002, China; Joint International Research Laboratory of Ecological Pest Control, Ministry of Education, Fuzhou 350002, China

**Keywords:** Conservation biological control, *Plutella xylostella*, natural enemy community, landscape composition, habitat diversity, multiple spatial scales, China

## Abstract

**Background:** Conservation biological control provides an environment-friendly approach to improve the efficacy of natural enemies. Although numerous studies have demonstrated the potential of semi-natural habitats in promoting biological control in organic or unsprayed agroecosystems, few studies were conducted in conventional agricultural fields. In this study, we investigated the effects of landscape composition on the major pests of cruciferous vegetables and on the assemblages of their natural enemies in southeastern China.

**Results:** Habitat diversity, particularly increasing grassland proportion in the landscape, had a positive impact in controlling both small-sized pests (aphids, leaf miners, thrips and flea beetles) and *Plutella xylostella*. This increasing proportion also promoted greater abundance and diversity of canopy-dwelling predators, more forests supported a higher diversity of airborne enemies (parasitoids and canopy-dwelling predators) as well as a higher abundance of ground-dwelling predators. A general increase in habitat diversity was beneficial to parasitoids and ground-dwelling predators. Additionally, the proportion of forest, grassland, and non-cruciferous vegetable area, as well as habitat diversity, affected the compositions of natural enemy communities. Moreover, inconsistent effects of non-cruciferous and grassland habitats were found between sampling regions for small-sized pests and canopy-dwelling predators. Moreover, the scale at which pests and natural enemies’ abundance and richness responded most to landscape composition varied with their feeding range and dispersal ability.

**Conclusion:** Our study provides evidence that increasing the amount of semi-natural habitats and habitat diversity can result in lower pest and higher natural enemy abundance in conventional cruciferous agroecosystems. Regional conditions and spatial scales also should be considered in designing the agricultural landscape mosaic.

## 1. Introduction

Conservation biological control (CBC) provides an alternative approach to conventional chemical-based farming, which allows to minimize negative effects, such as environment contamination and insecticide resistance, and also reduce costs. It emphasizes the importance of improving the efficiency of natural enemies by applying strict habitat regulations and reducing pesticide use (Eilenberg et al., 2001; Jonsson et al., 2008). Therefore, over the past decade, CBC has received a broad recognition and increasing research attention.

Previous studies have shown that semi-natural habitats (SNHs), such as forest, grassland, and hedgerow covered with perennial vegetation, are crucial for providing essential resources for natural enemies (Tscharntke et al., 2005), including nectar, pollen, shelter, and habitats for hibernation (Bianchi et al., 2006; Gurr et al., 2017). Moreover, increasing SNHs is generally associated with increases in the abundance and richness of natural enemies, thus with a high level of pest suppression (Grab et al. 2018). The positive effect on biological control is, however, not always achieved (Karp et al., 2018; Letourneau et al., 2009); negative, and neutral effects of enemy diversity could be caused by intraguild predation, and functional redundancy (Straub et al., 2008). Moreover, whilst different types of SNHs favor different natural enemies, increased proportion of some habitats can equally be beneficial for pests and their enemies (Holland et al., 2016). Nonetheless, landscape diversity can act as a driver of the structure of natural enemy assemblages in agricultural landscapes (Martin et al., 2016). Thus, investigating the composition of assemblages is key to understand agroecosystem function and, in turn, the effectiveness of biological control (Bommarco et al., 2013; Rusch et al., 2014).

Although there are numerous studies showing the benefits of SNHs, in these unsprayed or organic crop areas are overrepresented (Perez-Alvarez et al., 2018; Thies et al., 2011) and the effects of increased SNH proportions in conventional (high pesticide input) farming systems are rarely investigated. Although Madeira et al. (2016) showed that arthropod communities in conventional farms also heavily depend on the nearby source habitats (such as grassland), there is evidence that pest control promoted by increased SNH areas is more efficient, or even only possible, in unsprayed fields (Gagic et al., 2019; Saqib et al., 2020). Thus, uncertainty in the impact of management remains, and there is an urgent need to assess the benefits of SNHs for natural enemies in real-world settings with conventional management strategies.

Landscape composition variables may also affect pests and natural enemies at different spatial scales or even at multiple spatial scales simultaneously (Martin et al., 2016; Ortega and Pascual, 2014; Sivakoff et al., 2013). Indeed, Gonthier et al. (2014) suggested that preservation of multiple taxonomic groups in agriculture requires a multi-scale approach in conservation. However, studies investigating the effect of SNHs are often restricted to narrow spatial scales (Bailey et al., 2010; Concepción et al., 2008; Muneret et al., 2019b), and only to a small number of species (Hermann et al., 2013; Perović et al., 2010). Thus, our understanding on the effects of landscape composition on multiple pests and natural enemies at multiple spatial scales, particularly in conventional farms, is still limited. The lack of this knowledge may hamper the effective biological control in the broader landscape.

Cruciferous vegetables are often attacked simultaneously by multiple insect pests (e.g. aphids and flea beetles), whose impact can be influenced by landscape habitat diversity (Perez-Alvarez et al., 2018; Zaller et al., 2008). Among these pests, the diamondback moth (DBM), *Plutella xylostella* (Linnaeus, 1758) (Lepidoptera: Plutellidae), is the most serious threats to cruciferous vegetables production in China (Li et al., 2016). Although numerous studies have focused on the biology and ecology of this pest (Furlong et al. 2004; Furlong, Wright, and Dosdall 2013; Gurr et al. 2018), effective management is still limited. Considering landscape composition can be a potential way of increasing the efficiency of control effort.

Thus, to address knowledge gaps, we investigated (1) how landscape composition affects the abundance and assemblage structure of pests in cruciferous vegetable fields along different spatial scales, and (2) how landscape composition affects the abundance, richness, and assemblage structure of natural enemies along different spatial scales? Therefore, we selected 23 fields over a gradient of SNHs cover (including forests and grasslands), and sampled both pest and enemy insects over two years in three typical cruciferous vegetables planting regions in Fujian Province, southeastern China. We hypothesized that:

1. Increasing proportion of SHNs (e.g. forest and grassland) and landscape diversity have a positive effect on the abundance and richness of natural enemies, but a negative one on pests.
2. Landscape composition variables significantly affect the composition of natural enemies’ assemblages.
3. Scales of response to different landscape composition variables vary within different pest and natural enemy groups.

## 2. Material and methods

### 2.1 Study area

The study was conducted in cauliflower farms in Fujian Province, southeastern China (Figure 1A). A total of 23 fields were selected in three different locations, with 9 fields at a sampling location of Ningde Municipal Region, 7 fields at a location of Nanping Municipal Region, and 7 fields at a location of Zhangzhou Municipal Region. All fields were at least 1km apart each other, along a gradient of proportion of SHNs in the landscape. These locations represent the three most typical types of conventional vegetable planting regions. Ningde region is characterized by a succession cropping system in a mountainous area. Cruciferous vegetables are planted both in autumn and spring. Nanping and Zhangzhou regions are typical cruciferous-rice rotation cropping systems along a river with cruciferous vegetables planted only in autumn and rice is planted in spring. Local management was homogenous across fields in each region.

**Figure 1.**
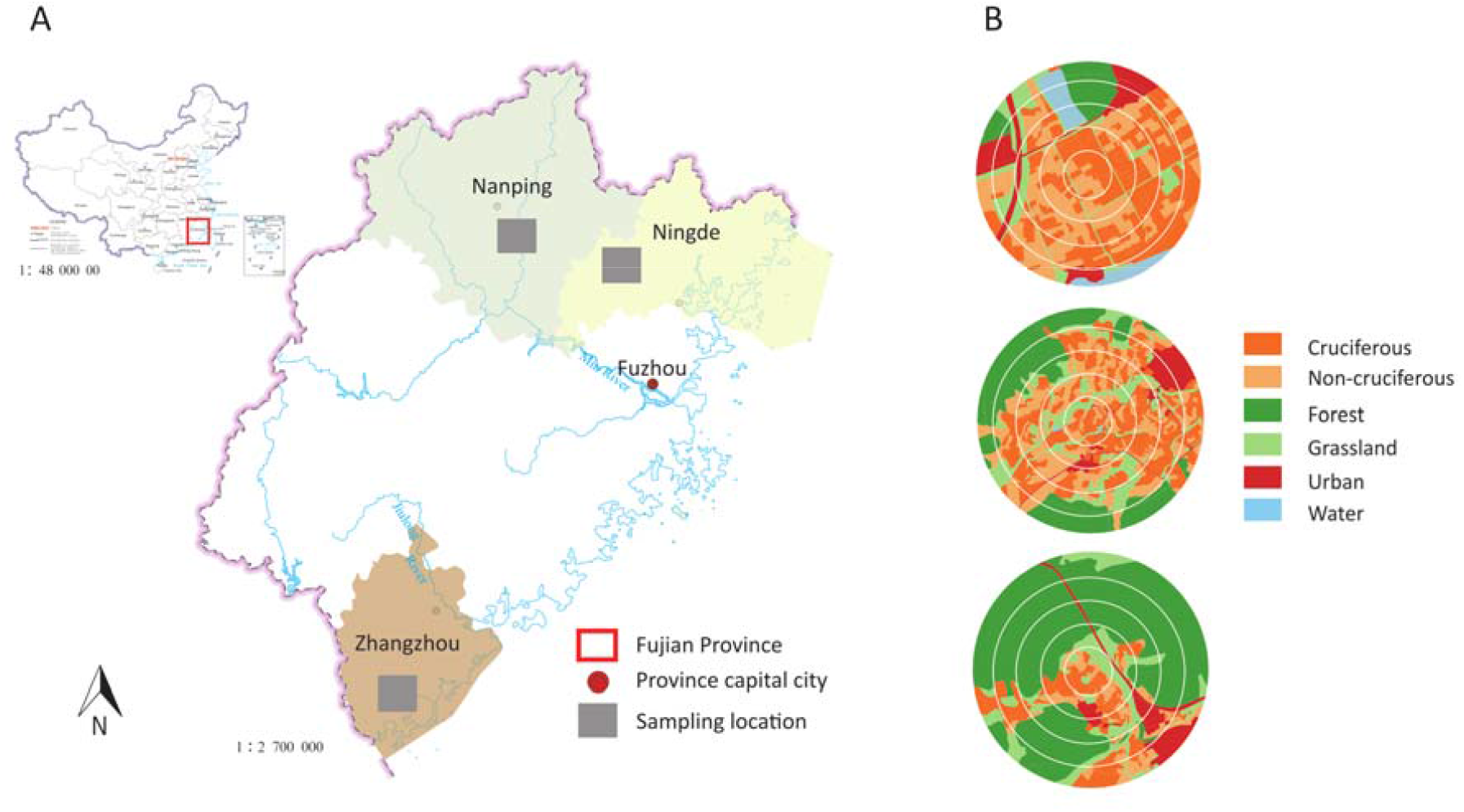
Map showing sampling locations in Nanping, Ningde, Zhangzhou of Fujian Province, southeastern China, and examples of mapping different habitats. (A) Locations of sampling locations in Fujian Province of China. (B) Examples of mapping different habitats within 100, 200, 300, 400 and 500 m radii buffers around focal patch.

In each field, a 20 × 20 m focal patch was chosen for sampling. Landscape compositions were quantified within a 500 m radius around the focal patch. The 500 m radius was selected because natural enemies have been found to respond strongly to landscape composition at spatial scales <500 m (Bianchi, Goedhart, and Baveco 2008; Djoudi et al. 2018; Jonsson et al. 2015).

### 2.2 Landscape analysis

To investigate the habitats distribution in each fields, we took aerial photos of each sampling fields by a drone (PHANTOM 4, Shenzhen Dajiang Baiwang Technology Co., Ltd., China) in a 500 m radius circle. The vegetation types were identified using the aerial photos and validated by field investigations. Habitats types in the 500 meter circle were categorized into grassland, forest, urban areas (e.g., residential land, greenhouses and roads), water surfaces (e.g. rivers, small steam, and ponds), cruciferous vegetables, and non-cruciferous vegetables (e.g. pepper, eggplant, corn, and beans) (Table S1). The proportion of different habitats types was calculated using QGIS 3.4. Landscape compositional diversity was measured using Shannon-Wiener diversity index (SHDI). Proportion of different habitats and SHDI was calculated at multiple spatial scales by splitting the 500 m radius landscape surrounding the focal patch to 5 concentric buffer circles in every 100m intervals (Figure 1B).

### 2.3 Arthropod sampling

Sampling was conducted in the autumn of 2017 and 2018, focusing mainly on pre-harvest period that is generally associated with a maximum abundance of arthropods.

#### Pests

Adult *P. xylostella* was sampled in every field for three times each year, using five boat-shape traps (Figure 2A) with sex pheromone (Pherobio Technology Co., Ltd, Beijing, China). The sticky papers were changed every five days, and the *P. xylostella* individuals were counted. Small-sized pests (including aphids, leaf miners, thrips and flea beetles) were sampled using yellow pan traps (Figure 2B), with a transparent plastic board installed onto each trap to improve capture rate. In each field, five boat traps and five pan traps at vegetable height were set in the focal patch (one in the center and four in the corner of square), each traps was set up 20 m apart.

**Figure 2.**
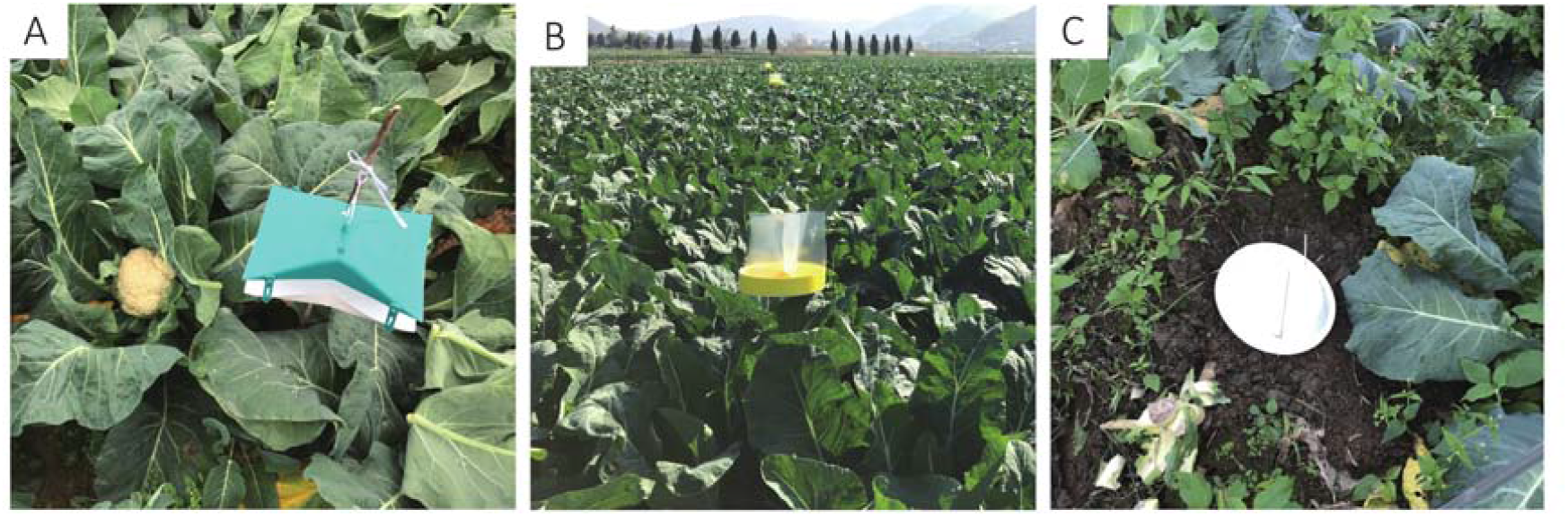
Sampling methods for pests and natural enemies. (A) Pheromone traps, (B) Yellow pan traps, and (C) Pitfall traps.

#### Natural enemies

Airborne enemies (including Vespidae, Reduviidae, ballooning spiders, and parasitoids) were also sampled by yellow pan traps. They were grouped into two functional groups: canopy-dwelling predators and parasitoids. Beside pan traps, five pitfall traps (Figure 2C) were set to sample ground-dwelling natural enemies (mainly spiders, carabids, and rove beetles) in each field. The pitfall traps were (9.5 cm diameter and 13.8 cm height) half-filled with propylene glycol solution with a few drops of added detergent to reduce surface tension. A 10 cm diameter roof above each pitfall traps was installed to prevent the rain diluting antifreeze. All specimens were transferred into 80% ethanol in tubes, and identified to species or morphospecies.

### 2.4 Sorting and identification

For each site, arthropods were sorted into groups of pests, parasitoids, and predators and identified to, at least, family level. This level of identification was confirmed to be a meaningful proxy for assessing species-level diversity patterns in biodiversity studies (Timms et al., 2013; Zou et al., 2020). The majority of parasitoids were identified to family level, but Braconidae and Ichneumonidae were further sorted by experts (See Acknowledgements) to subfamily level. Cynipidae and Agaonidae were excluded from the dataset because they do not parasitize insects. We first photographed predator individuals (e.g. Carabidae, Formicidae, Reduvidae and Araneae) and assigned them to a morphospecies, then, we used molecular identification to determine the actual species.

Genomic DNA of each assigned morphospecies was extracted from an excised leg from a representative specimen. Polymerase chain reaction (PCR) was performed to amplify the ∼658 bp region of mitochondrial COI gene using established universal primer pair for arthropods. The universal primer and reaction environment was the same in Saqib et al. (2020). Each amplified DNA of morphospecies was annotated based on similarity with available sequences in GeneBank (https://www.ncbi.nlm.nih.gov/) to identify the sorted morphospecies.

### 2.5 Statistical analysis

The relationship between the abundance and richness of pests and natural enemies and landscape composition was analyzed using a series of generalized linear mixed □effects models (GLMM). For individual models, response variables were (1) the abundance of *P. xylostella*, (2) the abundance of small-sized pests, (3) the abundance of parasitoids, (4) the number of families (henceforth family richness) of parasitoids, (5) the abundance of canopy-dwelling predators, (6) the family richness of canopy-dwelling predators, (7) the abundance of ground-dwelling predators, and (8) the family richness of ground-dwelling predators. As explanatory landscape composition variables, the proportion of urban area and water body were deleted from our final models because they were not present in every investigated region. Moreover, we only included habitats which were likely to support agroecologically important arthropods: the proportion of (a) cruciferous vegetables, (b) non-cruciferous vegetables, (c) forests, (d) grasslands, and (e) SHDI that was calculated based on all types of habitats. Sampling regions were included as interacting variables with landscape composition variables. Study field and sampling date were added as random variables. Variance inflation factor (VIF) was calculated to check for multicollinearity between explanatory variables, using the ‘car’ R package (Fox and Weisberg, 2019). As a result of its high collinearity with other habitat proportions (VIF>5) (Zuur et al., 2009), the proportion of cruciferous vegetables was excluded in our final models. Analyses were conducted at five spatial scales, taking the landscape composition in a radius of 100, 200, 300, 400 and 500 m around the focal patch.

Arthropod data of five traps in each study field at each sampling date were pooled prior to analyses. To ensure the use of the appropriate distribution type and link function were used, each model was checked for overdispersion. Poisson distribution was selected for richness of parasitoids, canopy-dwelling predators and ground-dwelling predators. Negative binomial distribution was used to model the abundance of *P. xylostella*, small-sized pests, parasitoids, canopy-dwelling predators and ground-dwelling predators.

All predictor variables were standardized (z□scored transformation) prior to analysis (Grueber et al., 2011) using the ‘standardize’ function of the R package ‘arm’ (v. 1.11-2; Gelman and Su, 2020). Then the ‘dredge’ function of the R package ‘MuMin’ (v. 1.43.1; Barton, 2020) was used to fit all possible combinations of models, and to calculate their associated corrected Akaike’s Information Criteria (AICc) (Liddle et al., 2009). The best fitted models with Δ AICc < 2 were obtained with the ‘get.models’ function and they were then averaged with the ‘model.avg’ function in the ‘MuMIn’ package.

To avoid the effect of rare species in analyses of community composition, only the five most abundant families of parasitoids, canopy-dwelling predators and ground-dwelling predators were analyzed. Redundancy analysis (RDA) was then used to detect the relationships between landscape variables and the composition of natural enemy assemblages. The community dataset was transformed to a Hellinger distance matrix prior to analyses (Borcard et al., 2011). We used landscape composition variables and regions as predictors. We also checked for multicollinearity between explanatory variables in the RDA model by calculating VIFs. Variables with high collinearity (VIF> 5) were removed from final models (Belsley et al., 2005). Thus, in the final model, proportion of cruciferous habitat type was excluded from the landscape variables. To assess the significance of RDA models, a permutation test with 999 iterations was carried out to assess the significance of RDA models and each environmental factors, using R package ‘vegan’ (Legendre et al., 2011).

## 3. Results

### 3.1 Community composition and abundance of pests and natural enemies

A total of 3,952 individuals of *P. xylostella* and 46,482 individuals of small-sized pests were captured, with the dominant pests being Aphididae (87.58%). Altogether, 11,888 individuals of natural enemies were caught, representing 7 orders, 53 families. Of the 4,068 individuals of parasitoids, Encyrtidae (22.28%), Braconidae (16.60%), Diapriidae (11.50%), Eulophidaes (9.41%) and Ichneumonidae (9.12%) were the dominant families. The 6,202 individuals of canopy-dwelling predators captured, Staphylinidae (72.62%) and Linyphiidae (14.38%) were the dominant families. Among the 1,618 individuals of ground-dwelling predators, Formcidae (33.99%) and Carabidae (24.17%) were the most abundant (Table S3). The dominance in relative abundance, however, varied among sampling regions (Figure 3).

**Figure 3.**
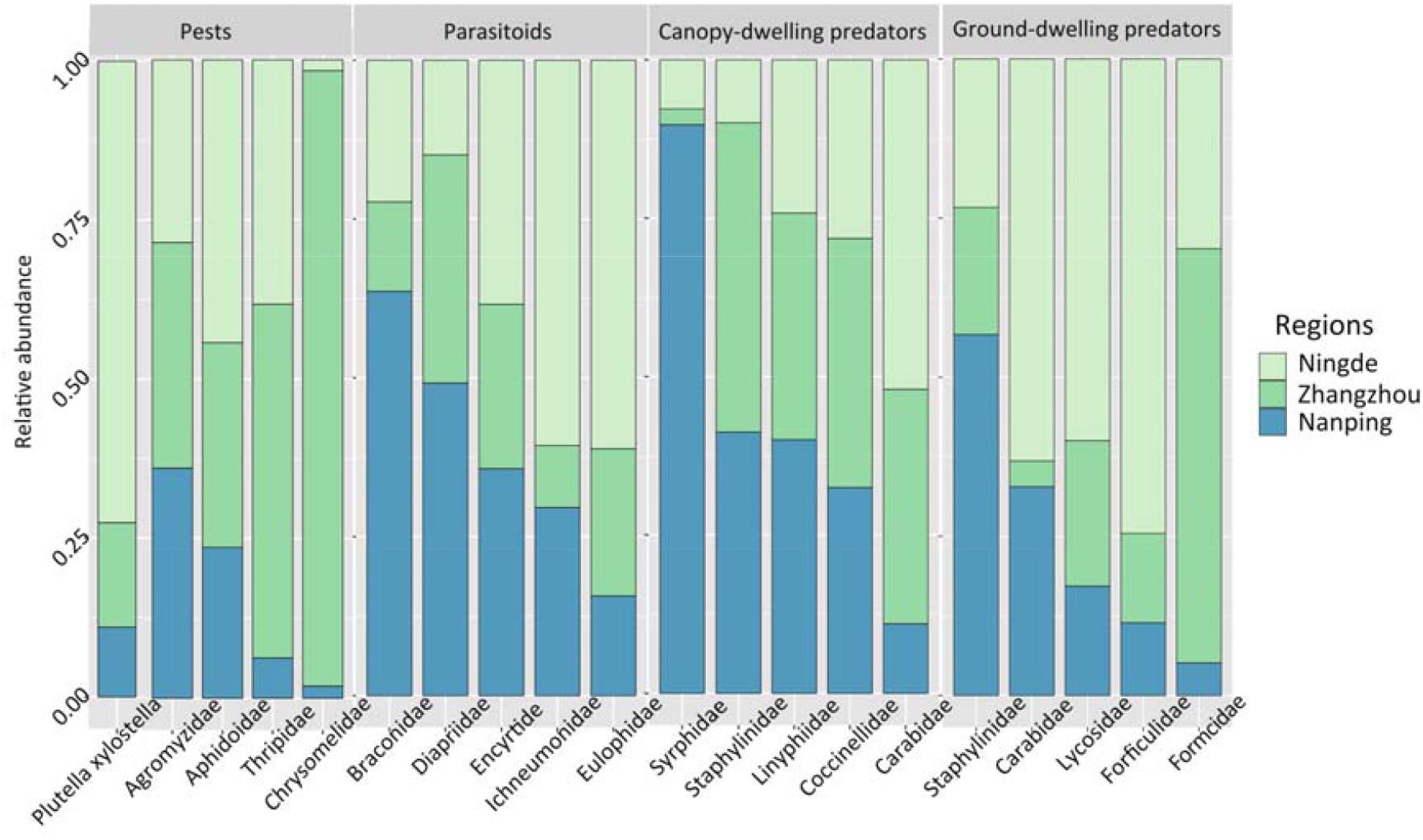
Relative abundance of pests and natural enemies in different sampling regions.

### 3.2 Effects of landscape variables on pests and natural enemy community structure

The GLMM showed that the abundance of *P. xylostella* was negatively associated with the proportion of grassland, but positively associated with the proportion of non-cruciferous vegetables. Similarly, the abundance of small-sized pests was found to be negatively associated with SHDI but positively associated with the proportion of forests (Figure 4). In addition, the proportion of forests improved the diversity of parasitoids and canopy-dwelling predators, and support more ground-dwelling predators. The proportion of grassland positively affected the richness of canopy-dwelling predators. Both parasitoids and ground-dwelling predators were found to be positively correlated with SHDI (Figure 4).

**Figure 4.**
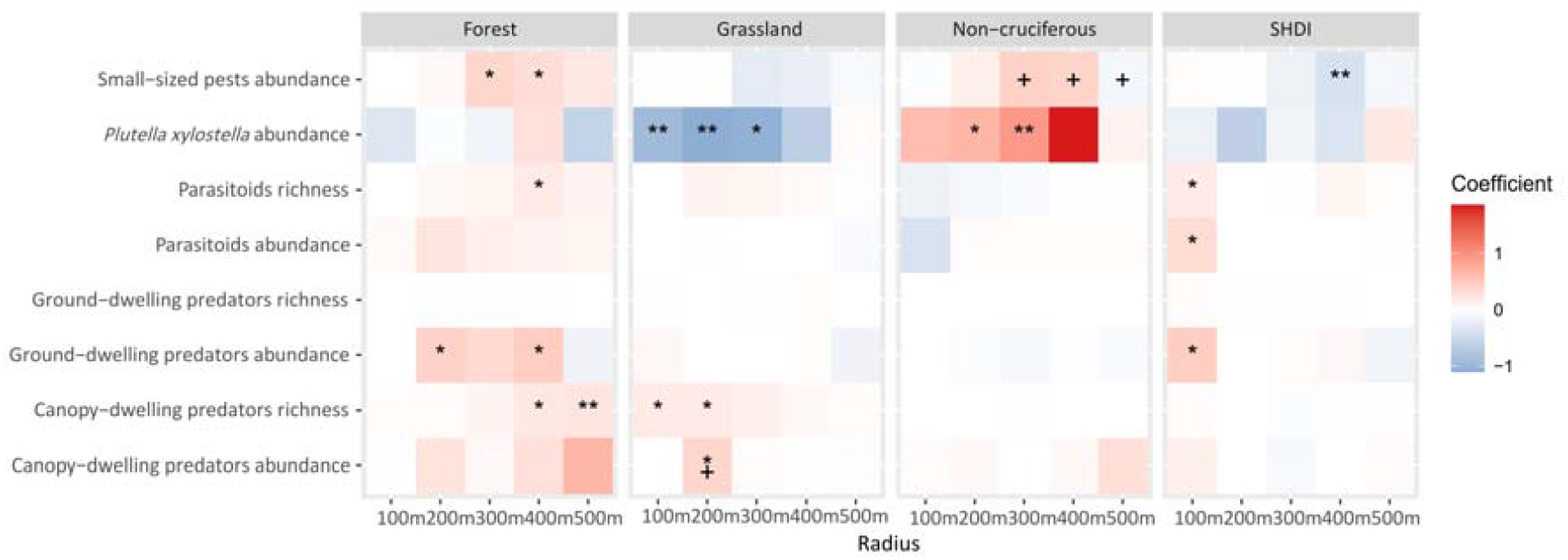
Relationship between the abundance/richness of *Plutella xylostella*, small-sized pests, parasitoids, canopy-dwelling predators, ground-dwelling predators and landscape composition variables at a 100, 200, 300, 400 and 500 m radius. The gradient of the color represents the value of the GLMM estimated coefficients. “*”indicates a significant correlation at *P* < 0.05 level, and”**” at *P* < 0.01 level. “**+**”indicates the interaction between landscape variables with sampling regions.

The proportion of non-cruciferous vegetables and grasslands showed inconsistent effects for small-sized pests and canopy-dwelling predators between sampling regions (Figure 4-5, Table S2).

**Figure 5.**
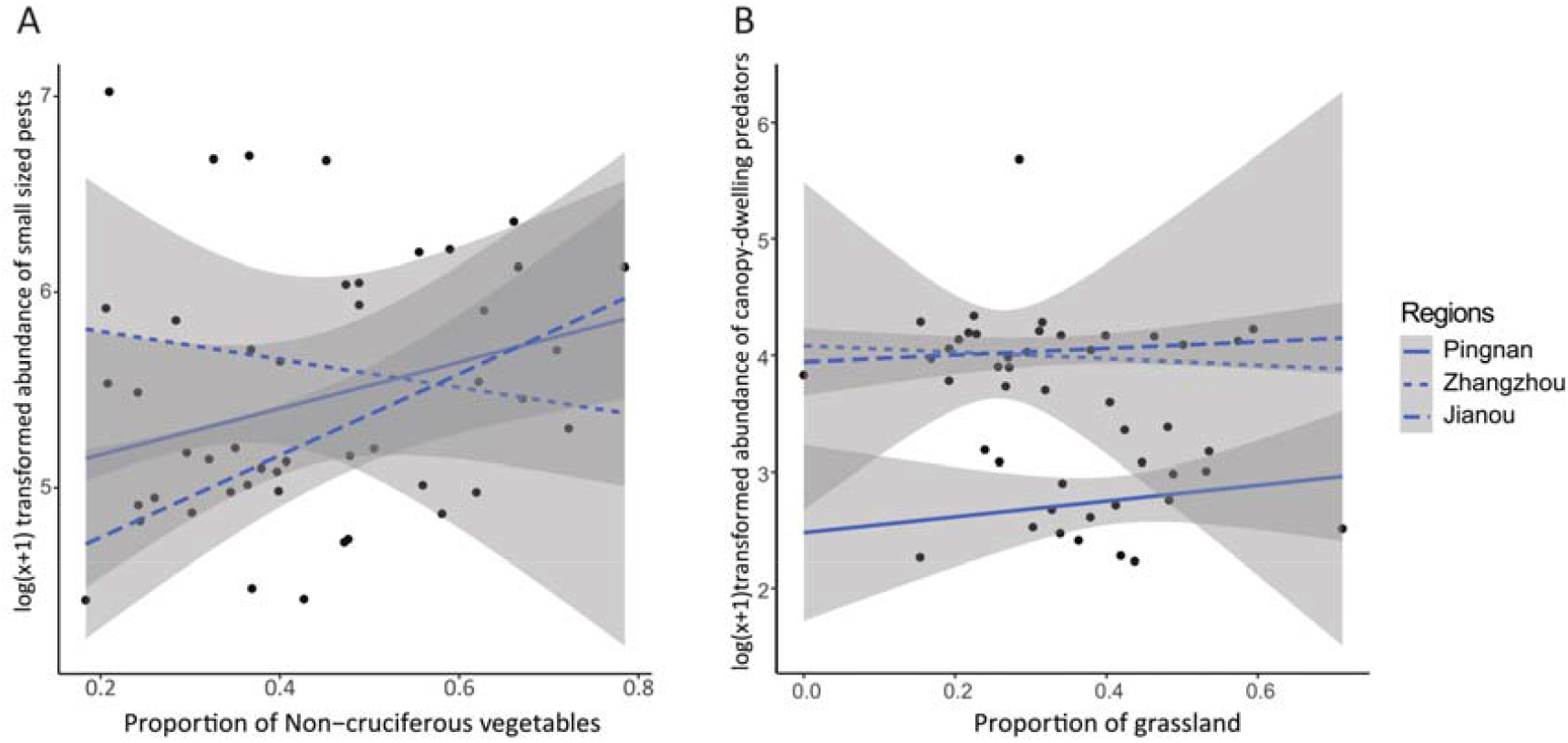
Effects of the (A) proportion of non-cruciferous vegetables on small-sized pests (at 400m), and (B) proportion of forests on the abundance of canopy-dwelling predators (at 200m). Separate lines indicate different sampling regions.

The scale at which pests and natural enemies responded best to landscape composition variables varied among functional groups. The responses scale to landscape variables of small-sized pests (300 - 400m) is larger than *P. xylostella* (100 - 300m). Similarly, airborne enemies (including parasitoids and canopy-dwelling predators) showed response on a larger scale to forest proportion than ground-dwelling enemies (Figure 4).

### 3.3 Effects of landscape variables on natural enemy assemblage composition

The final RDA models explained 25.8% of the total variability in the assemblage structure of airborne enemies (including parasitoids and canopy-dwelling predators) and 19.2% of the total variability in that of ground-dwelling enemies. The results showed that sampling regions, proportion of non-cruciferous vegetables, and forests in the landscape cumulatively explained 97.9% variance of the assemblages of airborne enemies and 98.3% in that of the 5 most abundant families of ground-dwelling predators. Conversely, only a small fraction of variability of enemy assemblage was explained by the spatial scales (i.e. radius around the focal point). The assemblage structure of parasitoids, canopy-dwelling predators, and ground-dwelling predators were significantly influenced by sampling regions, non-cruciferous proportion, forest proportion, grassland proportion and SHDI in the agriculture landscape (Table 1).

**Table 1.** Permutation tests for redundancy analysis (RDA) of different groups of enemies in different regions with varying landscape composition variables.

| Factors | Parasitoids and canopy-dwelling predators |  | Ground-dwelling predators |  |
| --- | --- | --- | --- | --- |
|  | ChiSquare | Pr(>F) | ChiSquare | Pr(>F) |
| Regions | 0.054 | 0.001*** | 0.071 | 0.001*** |
| Radius | 0.000 | 1.000 | 0.000 | 1.000 |
| Non-cruciferous | 0.003 | 0.001*** | 0.012 | 0.001*** |
| Forest | 0.001 | 0.022* | 0.009 | 0.001*** |
| Grassland | 0.001 | 0.001*** | 0.003 | 0.005*** |
| SHDI | 0.001 | 0.001*** | 0.001 | 0.086 |
| RDA model | 0.061 | 0.001*** | 0.097 | 0.001*** |

“\*” indicates the significance when $P < 0.05$ , “\*\*” when $P < 0.01$ , and “\*\*\*” when $P < 0.001$ .

For the airborne enemies, fields with high proportion of forests were characterized by high abundance of species belonging to parasitoids (Ichneumonidae, Eulophidae and Encyrtidae) and those Carabidae species which are good at flying, but negatively correlated with Staphylinidae. Fields with high proportion of grassland favored more Braconidae and canopy-dwelling predators, such as, Linyphidae and Syrphidae. Non-cruciferous vegetables supported similar enemy assemblages with grassland. Fields with diverse habitats were characterized by high abundance of species belonging to Diapriidae and Carabidae (Figure 6A). For the ground-dwelling predators, high proportion of forests supported Staphylinidae, grasslands were positively correlated with Carabidae and Forficulidae, but negatively correlated with Formicidae (Figure 6B).

**Figure 6.**
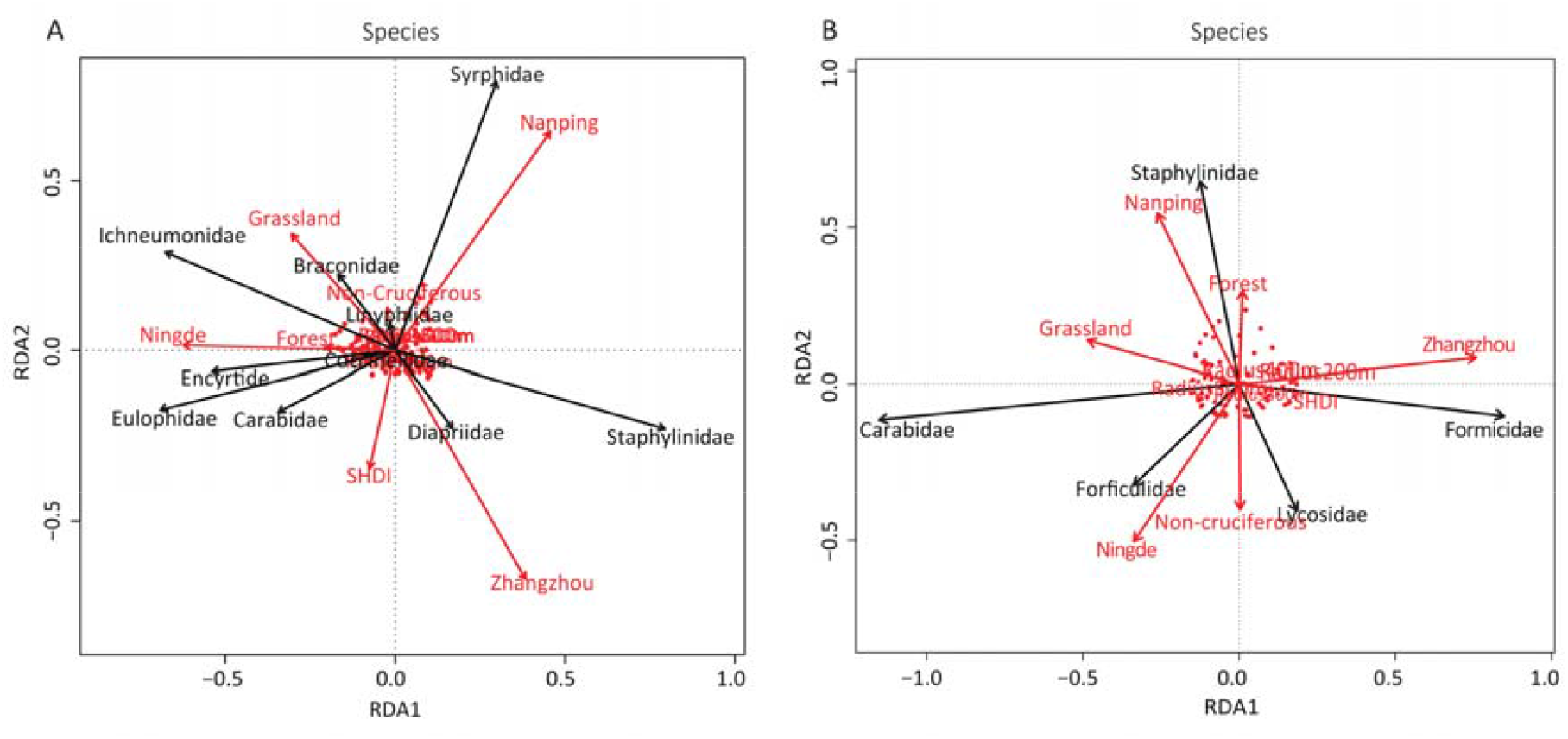
RDA ordination diagram represents the association of (A) Abundance of parasitoids and canopy-dwelling predators, and (B) Abundance of ground-dwelling predators in different regions across varying proportions of habitats.

## 4. Discussion

In this study we provided novel insights into the effects of landscape composition along different spatial scales on pests and different groups of natural enemies in conventional cruciferous vegetable fields. Similarly to previous studies (Dominik et al., 2018; Martin et al., 2016; Muneret et al., 2019a), the responses of natural enemy assemblages to landscape compositions differed from one functional group to another.

In agreement with our Hypothesis 1, we found a negative relationship between *P. xylostella* and the proportion of grasslands in the landscape. Although some studies demonstrated positive effects of non-crop habitats on parasitoids of lepidopteran pests in cruciferous fields (Bianchi, Goedhart, and Baveco 2008; Letourneau, Bothwell Allen, and Stireman 2012; Perez-Alvarez, Nault, and Poveda 2018), we did not find significantly increased parasitoid abundances with the increasing proportion of grasslands. High proportion of grasslands supported higher abundance and diversity of canopy-dwelling predators though, suggesting that predators play the most important role in controlling *P. xylostella* in cruciferous vegetables. Some work showed that pests (e.g. flea beetles and aphids) also positively correlate with the proportion of grasslands (Perez-Alvarez et al., 2018), but in our study, the lack of positive relationship between small-sized pests (mainly aphids) and grassland area did not provide further support. On the other hand, in line with Zaller et al. (2008), we demonstrated that increasing forests area, but not grassland, promoted small-sized pests’ abundance. Thus, although forests promoted diversity of airborne enemies and abundance of ground enemies, their effect in suppressing small-sized pests is controversial.

The reason could be the small-sized pests populations were more driven by environment condition or agriculture practices than predation or parasitism (Tscharntke et al., 2016). Unexpectedly, the area proportion of non-cruciferous vegetables showed significantly positively effects on *P. xylostella*. None of the natural enemy groups were found to be positively influenced by non-cruciferous vegetables. Here, agriculture practices can offset naturally occurring “top-down” control (Bommarco et al., 2011; Tscharntke et al., 2016), and the positive effect of biocontrol agents is less likely. Moreover, since non-cruciferous vegetables were also heavily managed, the higher number of *P. xylostella* compared to that of natural enemies may be a result of its higher tolerance to management, particularly to pesticide use (Furlong et al., 2013). Similarly to Dominik et al. (2018), increasing landscape diversity decreased the abundance of small-sized pests. Since in our case increasing landscape diversity most often increase the area of SHNs (e.g. forest and grass) that promote diversity predators and parasitoids, the general decline in pest numbers along with increased landscape diversity is not surprising. In diverse landscapes, a mosaic of cultivated and natural habitats provide a continuous supply of resources for predators and parasitoids over space and time, helping them to avoid spatial and temporal disturbance (Schoenly et al., 2010).

Our results reveal that whilst predators can benefit from grasslands and facilitate the suppression of *P. xylostella*, forests provide benefits to both small-sized pests and multiple natural enemies. However, the benefit of increasing some SNHs may come with some trade-offs. Perez-Alvarez et al. (2018) showed that high proportion of grasslands were associated with lower densities of *Pieris rapae* (Linnaeus, 1758) (Lepidoptera: Pieridae), but also with greater pest pressure by aphids and flea beetles. Similarly, in our study, increasing the proportion of grassland around cruciferous fields negatively affected lepidopteran pests (e.g. *P. xylostella*), but the increase of aphids and flea beetles were not pronounced. Forest, on the other hand, supported aphids and flea beetles in greater abundances. One explanation can be that high intensity management practices (e.g. insecticides or fungicides) disturbed the grassland environment adjacent to vegetables fields and forced these small bodied pests to seek refuge farther. Alternatively, we cannot exclude a basic host preference to the plant species living in woodlands. Nevertheless, there may be a trade-off between increasing the proportion of forests to enhance biological control and an increase in damage by small-sized pest.

In agreement with our Hypothesis 2, our results also showed that landscape composition variables can markedly affect the assemblage composition of natural enemies. In landscapes where the forest was dominant, canopy-dwelling predators (e.g. Carabidiae *Stenolophus mixtus* (Herbst, 1784)) and ground-dwelling Staphylinidae (e.g. *Quedius capucinus* (Gravenhorst, 1806)) with high dispersal ability and parasitoids were abundant. This was likely due to the stable understory environment with the high diversity of plant species and alternative prey (González et al., 2020). In landscapes with high proportion of grasslands, Braconidae and Syrphidae that are specialist enemies linked to open areas, and omnivorous predators like Linyphiidae (e.g. *Erigone prominens* Bösenberg & Strand, 1906), ground-dwelling Carabidae (e.g. *Trichotichnus longitarsis* A.Morawitz, 1863 and *Harpalus calceatus* (Duftschmid, 1812)) and Forficulidae were more abundant. In terms of dispersal ability, these predators are weaker than those in forests. Non-cruciferous vegetables also played a role in driving the community composition of enemies. Though they support similar airborne enemy assemblages as grasslands, the supported assemblages of ground-dwelling enemies are different. Most commonly, these enemies were generalist predators with high mobility that allows them both to avoid the frequent disturbance, and simultaneously exploit abundant prey resources in vegetable areas. For instance, Lycosidae with limited dispersal ability but high mobility are likely to move shorter distances between adjacent vegetable fields but not from farther grasslands or forests.

In our study, inconsistent effects of non-cruciferous and grassland habitats were found between sampling regions for small-sized pests and canopy-dwelling predators. This inconstancy can be due to different local managements, such as rotation cropping and farming intensity, or environment conditions between regions (Karp et al., 2018; Martin et al., 2016). Indeed, studies demonstrated that regional differences are important factors in shaping arthropod assemblages (Dominik et al., 2017; Massaloux et al., 2020; Schweiger et al., 2005), and thus, these also need to be considered when sustainable agricultural landscapes, to provide high level of biological control, are designed.

In agreement with our Hypothesis 3, the scales at which pests and natural enemies most responded to landscape compositions varied among functional groups and reflected their feeding range and dispersal abilities (Chaplin-Kramer et al., 2011; Dominik et al., 2018; Martin et al., 2016). The scale of responses to landscape variables of small-sized pests (300 m to 400 m) was larger than that of *P. xylostella* (100 m to 300 m). These results are in line with previous studies showing larger response scales for generalist than specialist pests (Chaplin-Kramer et al., 2011). Ground-dwelling predators are often wingless (e.g. Lycosidae) or limited in flight (e.g. Forficulidae) and are likely to move over shorter distances in vegetable fields. Thus, the scale at which they responded (200 and 400 m) to forests was found to be lower than highly mobile airborne enemies such as parasitoids (400 m) and canopy-dwelling predators (400 and 500 m). The richness of parasitoids was better predicted on a smaller scale than that of canopy-dwelling predators (e.g. Syrphidae or Vespidae), presumably because parasitoids’ smaller body size negatively correlated with their dispersal distance (Martin et al., 2016; Ritchie and Olff, 1999). Those taxa with high flight capabilities (such as Syrphidae) may respond on a larger than the 500 m scale we investigated (Martin et al., 2016), and thus, their dependence on landscape complexity can be masked in our results.

## 5. Conclusions

Our study shows the effects of different landscape composition variables on shaping arthropod communities in conventional cruciferous agroecosystems. We provide evidences that increasing proportion of semi-natural habitats and Shannon diversity in the landscape surrounding vegetable fields can result in lower pest abundance and more natural enemies. This effect is not clear though, in our study only forest and grassland areas showed clear positive effects on natural enemies and even these varied with spatial scales. Thus, maintaining or even improving the diversity of agricultural and semi-natural landscape mosaics has the potential to enhance the “top-down effects” on pests suppression (Gurr et al., 2017), but a case-by-case assessment may be needed to minimize trade-offs and maximize net benefits.

Based on these findings, we believe that both policymakers and stakeholders should aim for developing and implementing biodiversity improvement strategies to promote biological control in cruciferous agroecosystems, even in conventional farms. These strategies should include landscape management practices that increase landscape diversity at least 100–400 m around the fields, and particularly focus on improving natural forests and grasslands. This approach would be particularly beneficial for Chinese smallholders whose high pesticide use threatens healthy environment and is not line with global sustainability goals. Beside pest suppression, diverse landscape have other additional ecosystem services for conservation, such as, high biodiversity, pollination, ecological corridors, natural beauty, and human health (e.g. aesthetic values and mental health) (Fiedler et al., 2008; Vialatte et al., 2019).

## Acknowledgements

We thank all the farmers for their permission to sample their vegetable fields. We also thank Tao Li, Lingfei Peng, and Jun Li to help identify the parasitoids samples. This work was supported by the National Key R&D Program of China [2017YFD0200400] and the State Key Laboratory of Ecological Pest Control for Fujian and Taiwan Crops [2020L3007].

## Authors Contributions

JZ, GP, SY and MY conceived the ideas and design of the study and led the writing of the manuscript. JZ, DN, GGKG, AW and YY conducted fieldwork. JZ, GP and HSAS analyzed the data. All authors revised the final version and gave their approval for submission.

