## Supplementary material for "Landscape composition mediates suppression of major pests by natural enemies in conventional cruciferous vegetables": Table S1

**Table S1. Mean and range of habitats (%) within 100, 200, 300, 400 and 500 m radius in 2017 and 2018**

| Year | Regions | Radius | Habitats | | | | | |
| --- | --- | --- | --- | --- | --- | --- | --- | --- |
|  |  |  | Cruciferous | Non-cruciferous | Forest | Grassland | Urban | Water |
| 2017 | Ningde City | 500m | 0.14(0.02-0.26) | 0.24(0.06-0.33) | 0.48(0.22-0.79) | 0.07(0.03-0.12) | 0.06(0.01-0.17) | 0.00(0.00-0.00) |
| 2017 | Ningde City | 400m | 0.17(0.04-0.31) | 0.25(0.08-0.50) | 0.44(0.14-0.76) | 0.09(0.07-0.19) | 0.06(0.01-0.14) | 0.00(0.00-0.00) |
| 2017 | Ningde City | 300m | 0.19(0.06-0.32) | 0.27(0.07-0.58) | 0.38(0.05-0.70) | 0.11(0.07-0.22) | 0.06(0.03-0.10) | 0.00(0.00-0.00) |
| 2017 | Ningde City | 200m | 0.23(0.11-0.30) | 0.29(0.06-0.58) | 0.31(0.01-0.61) | 0.13(0.02-0.26) | 0.04(0.00-0.10) | 0.00(0.00-0.00) |
| 2017 | Ningde City | 100m | 0.34(0.22-0.56) | 0.33(0.06-0.54) | 0.17(0.00-0.35) | 0.16(0.00-0.31) | 0.00(0.00-0.02) | 0.00(0.00-0.00) |
| 2017 | Zhangzhou City | 500m | 0.26(0.12-0.43) | 0.18(0.05-0.43) | 0.29(0.07-0.56) | 0.06(0.05-0.09) | 0.14(0.01-0.35) | 0.07(0.01-0.30) |
| 2017 | Zhangzhou City | 400m | 0.28(0.12-0.51) | 0.18(0.04-0.44) | 0.27(0.07-0.54) | 0.06(0.04-0.08) | 0.14(0.00-0.36) | 0.06(0.01-0.22) |
| 2017 | Zhangzhou City | 300m | 0.33(0.13-0.65) | 0.18(0.05-0.42) | 0.27(0.1-0.52) | 0.06(0.04-0.12) | 0.12(0.00-0.29) | 0.04(0.01-0.09) |
| 2017 | Zhangzhou City | 200m | 0.40(0.17-0.72) | 0.15(0.07-0.32) | 0.28(0.08-0.47) | 0.07(0.03-0.17) | 0.06(0.00-0.13) | 0.04(0.00-0.10) |
| 2017 | Zhangzhou City | 100m | 0.52(0.24-0.85) | 0.10(0.02-0.24) | 0.30(0.05-0.56) | 0.05(0-0.14) | 0.02(0.00-0.11) | 0.02(0.00-0.10) |
| 2017 | Nanping City | 500m | 0.2(0.08-0.39) | 0.23(0.11-0.39) | 0.36(0.06-0.66) | 0.14(0.1-0.23) | 0.06(0.00-0.15) | 0.00(0.00-0.00) |
| 2017 | Nanping City | 400m | 0.25(0.1-0.46) | 0.25(0.13-0.42) | 0.31(0.04-0.59) | 0.14(0.1-0.22) | 0.05(0.00-0.15) | 0.00(0.00-0.00) |
| 2017 | Nanping City | 300m | 0.3(0.14-0.55) | 0.28(0.13-0.51) | 0.23(0.01-0.46) | 0.16(0.06-0.26) | 0.03(0.00-0.12) | 0.00(0.00-0.00) |
| 2017 | Nanping City | 200m | 0.38(0.15-0.78) | 0.29(0.09-0.61) | 0.18(0.00-0.34) | 0.13(0.00-0.31) | 0.03(0.00-0.17) | 0.00(0.00-0.00) |
| 2017 | Nanping City | 100m | 0.53(0.27-0.86) | 0.27(0.01-0.66) | 0.07(0.00-0.32) | 0.11(0.00-0.38) | 0.02(0.00-0.13) | 0.00(0.00-0.00) |
| 2018 | Ningde City | 500m | 0.17(0.08-0.34) | 0.18(0.04-0.67) | 0.47(0.11-0.77) | 0.12(0.00-0.20) | 0.05(0.01-0.16) | 0.00(0.00-0.01) |
| 2018 | Ningde City | 400m | 0.20(0.09-0.38) | 0.13(0.04-0.3) | 0.46(0.09-0.74) | 0.15(0.11-0.26) | 0.06(0.02-0.15) | 0.00(0.00-0.01) |
| 2018 | Ningde City | 300m | 0.22(0.11-0.39) | 0.16(0.06-0.34) | 0.37(0.02-0.67) | 0.18(0.14-0.33) | 0.06(0.01-0.14) | 0.00(0.00-0.01) |
| 2018 | Ningde City | 200m | 0.23(0.01-0.40) | 0.24(0.09-0.6) | 0.26(0.05-0.58) | 0.21(0.11-0.43) | 0.05(0.00-0.11) | 0.01(0.00-0.02) |
| 2018 | Ningde City | 100m | 0.41(0.21-0.61) | 0.25(0.18-0.4) | 0.10(0.00-0.35) | 0.22(0.07-0.39) | 0.02(0.00-0.09) | 0.00(0.00-0.01) |
| 2018 | Zhangzhou City | 500m | 0.27(0.11-0.53) | 0.15(0.03-0.31) | 0.3(0.07-0.53) | 0.07(0.05-0.11) | 0.15(0.01-0.36) | 0.07(0.01-0.25) |
| 2018 | Zhangzhou City | 400m | 0.29(0.14-0.51) | 0.16(0.04-0.34) | 0.28(0.07-0.52) | 0.07(0.05-0.11) | 0.13(0.00-0.36) | 0.06(0.01-0.22) |
| 2018 | Zhangzhou City | 300m | 0.33(0.18-0.62) | 0.16(0.04-0.35) | 0.28(0.1-0.51) | 0.08(0.05-0.13) | 0.11(0.00-0.28) | 0.05(0.01-0.09) |
| 2018 | Zhangzhou City | 200m | 0.39(0.22-0.69) | 0.15(0.04-0.3) | 0.28(0.08-0.47) | 0.07(0.04-0.15) | 0.06(0.00-0.15) | 0.04(0.00-0.10) |
| 2018 | Zhangzhou City | 100m | 0.49(0.23-0.87) | 0.09(0.00-0.25) | 0.31(0.05-0.56) | 0.05(0.00-0.10) | 0.04(0.00-0.20) | 0.02(0.00-0.06) |
| 2018 | Nanping City | 500m | 0.21(0.08-0.38) | 0.16(0.02-0.35) | 0.39(0.07-0.68) | 0.13(0.10-0.16) | 0.08(0.00-0.18) | 0.03(0.00-0.08) |
| 2018 | Nanping City | 400m | 0.24(0.09-0.43) | 0.19(0.03-0.39) | 0.34(0.05-0.64) | 0.13(0.10-0.19) | 0.07(0.00-0.20) | 0.02(0.00-0.10) |
| 2018 | Nanping City | 300m | 0.29(0.11-0.48) | 0.23(0.05-0.48) | 0.26(0.02-0.52) | 0.15(0.07-0.21) | 0.05(0.00-0.18) | 0.02(0.00-0.12) |
| 2018 | Nanping City | 200m | 0.36(0.13-0.64) | 0.25(0.07-0.56) | 0.19(0.00-0.37) | 0.14(0.05-0.29) | 0.04(0.00-0.16) | 0.03(0.00-0.17) |

Non-cruciferous vegetables including eggplant, pepper, tomatoe, bean, and corn.

Forests including Coniferous, Bamboo and Broad leaved trees.

Grassland including the strip and patch grassland in cultivated area and natural grass between forest and cultivated land.

Urban including residential district, road and artificial greenhouse.

Water including river, pond and reservoir.
