## Supplementary material for "Landscape composition mediates suppression of major pests by natural enemies in conventional cruciferous vegetables": Table S2

**Table S2. Coefficients and SE from model averaging showing the relationship between abundance and richness of pests and natural enemies, and landscape variables and interactions with sampling regions at a 100, 200, 300, 400 and 500 m radius. A plus (+) indicates that a significant or interaction with sampling regions, and asterisks (*) show significance levels (*≤0.05; **≤0.01; ***<0.001).**

| Response | Scale  (m) | AICc | Intercept | Region | Non-cruciferous | Forest | Grassland | SHDI | Interaction with sampling regions | | | |
| --- | --- | --- | --- | --- | --- | --- | --- | --- | --- | --- | --- | --- |
|  |  |  |  |  |  |  |  |  | Non-cruciferous | Forest | Grass | SHDI |
| Small-sized pests abundance | 100 | 1683.0 | 5.4±0.17*** | - | -0.03±0.09 | - | 0.01±0.07 | 0.02±0.07 | - | - | - | - |
|  | 200 | 1682.9 | 5.45±0.24*** | - | 0.14±0.19 | 0.06±0.13 | -0.01±0.06 | -0.01±0.23 | - | - | - | - |
|  | 300 | 1678.6 | 5.43±0.26*** | - | 0.42±0.27 | **0.38±0.16*** | -0.27±0.21 | -0.19±0.17 | **+** | - | - | - |
|  | 400 | 1673.9 | 5.37±0.26*** | - | 0.39±0.25 | **0.31±0.15*** | -0.21±0.20 | **-0.39±0.15**** | **+** | - | - | - |
|  | 500 | 1679.9 | 5.40±0.28*** | - | -0.12±0.26 | 0.21±0.20 | -0.08±0.16 | -0.11±0.17 | **+** | - | - | - |
| *Plutella xylostella* abundance | 100 | 1005.7 | 3.08±0.33*** | **+** | 0.61±0.37 | -0.33±0.39 | **-0.93±0.34**** | -0.19±0.31 | - | - | - | - |
|  | 200 | 1000.7 | 3.26±0.30*** | **+** | **0.68±0.26*** | -0.03±0.13 | **-1.11±0.34**** | -0.64±0.79 | - | - | - | - |
|  | 300 | 1001.1 | 3.16±0.28*** | **+** | **0.93±0.31**** | -0.14±0.27 | **-1.07±0.46*** | -0.14±0.26 | - | - | - | - |
|  | 400 | 1001.3 | 2.96±0.39*** | **+** | 1.85±1.31 | 0.28±1.26 | -0.65±0.52 | -0.35±0.61 | - | - | - | - |
|  | 500 | 1014.0 | 3.13±0.32*** | **+** | 0.12±0.24 | -0.58±0.37 | 0.04±0.18 | 0.21±0.35 | - | - | - | - |
| Parasitoids  abundance | 100 | 1069.2 | 3.15±0.27*** | - | -0.38±0.41 | 0.05±0.12 | - | **0.32±0.15*** | - | - | - | - |
|  | 200 | 1073.0 | 3.05±0.18*** | - | 0.04±0.11 | 0.24±0.18 | -0.01±0.07 | - | - | - | - | - |
|  | 300 | 1074.3 | 3.05±0.19*** | - | 0.04±0.11 | 0.14±0.17 | - | - | - | - | - | - |
|  | 400 | 1074.3 | 3.05±0.18*** | - | 0.04±0.11 | 0.12±0.16 | - | - | - | - | - | - |
|  | 500 | 1074.3 | 3.05±0.18*** | - | 0.03±0.09 | 0.09±0.15 | -0.06±0.14 | -0.01±0.06 | - | - | - | - |
| Parasitoids  richness | 100 | 660.3 | 2.11±0.12*** | - | -0.18±0.16 | - | - | **0.18±0.07*** | - | - | - | - |
|  | 200 | 664.1 | 2.06±0.13*** | - | -0.11±0.14 | 0.06±0.10 | 0.10±0.13 | **-** | - | - | - | - |
|  | 300 | 665.9 | 2.03±0.11*** | - | -0.06±0.13 | 0.09±0.09 | 0.09±0.11 | 0.01±0.04 | - | - | - | - |
|  | 400 | 665.4 | 1.99±0.09*** | - | -0.01±0.07 | **0.19±0.09*** | 0.05±0.10 | 0.09±0.11 | - | - | - | - |
|  | 500 | 667.9 | 2.01±0.10*** | - | 0.01±0.05 | 0.10±0.10 | -0.02±0.06 | 0.02±0.06 | - | - | - | - |
| Canopy-dwelling predators abundance | 100 | 1149.0 | 2.70±0.16*** | **+** | 0.04±0.16 | -0.01±0.07 | **-** | 0.14±0.34 | - | **-** | - | - |
|  | 200 | 1146.3 | 2.60±0.16*** | **+** | 0.06±0.11 | 0.25±0.21 | **0.38±0.17*** | - | - | **+** | - | - |
|  | 300 | 1149.4 | 2.71±0.15*** | **+** | - | 0.06±0.14 | 0.04±0.11 | -0.08±0.20 | - | - | - | - |
|  | 400 | 1149.4 | 2.67±0.16*** | **+** | 0.06±0.28 | 0.28±0.36 | 0.01±0.10 | - | - | - | - | - |
|  | 500 | 1148.1 | 2.58±0.19*** | **+** | 0.29±0.45 | 0.67±0.56 | -0.02±0.17 | 0.04±0.11 | - | - | - | - |
| Canopy-dwelling predators richness | 100 | 508.2 | 1.38±0.04*** | - | - | 0.02±0.05 | **0.19±0.09*** | 0.03±0.07 | - | - | - | - |
|  | 200 | 507.7 | 1.38±0.04*** | - | - | 0.04±0.08 | **0.19±0.09*** | - | - | - | - | - |
|  | 300 | 506.9 | 1.38±0.04*** | - | -0.01±0.04 | 0.10±0.11 | 0.14±0.11 | -0.03±0.07 | - | - | - | - |
|  | 400 | 506.3 | 1.38±0.04*** | - | - | **0.21±0.09*** | 0.06±0.08 | -0.02±0.06 | - | - | - | - |
|  | 500 | 505.1 | 1.38±0.04*** | - | - | **0.24±0.09**** | 0.03±0.07 | -0.01±0.05 | - | - | - | - |
| Ground-dwelling predators abundance | 100 | 798.5 | 2.29**±**0.12*** | - | - | - | 0.07±0.16 | **0.45±0.20*** | - | - | - | - |
|  | 200 | 799.1 | 2.29±0.13*** | - | -0.04±0.12 | **0.41±0.19*** | - | - | - | - | - | - |
|  | 300 | 798.9 | 2.29±0.13*** | - | -0.09±0.18 | 0.34±0.24 | - | 0.02±0.10 | - | - | - | - |
|  | 400 | 798.7 | 2.29±0.13*** | - | -0.02±0.11 | **0.46±0.21*** | 0.02±0.09 | 0.07±0.17 | - | - | - | - |
|  | 500 | 801.4 | 2.38±0.21*** | - | -0.06±0.17 | -0.16±0.38 | -0.16±0.23 | -0.14±0.24 | - | - | - | - |
| Ground-dwelling predators richness | 100 | 412.2 | 1.19±0.05*** | - | - | - | 0.01±0.05 | 0.02±0.06 | - | - | - | - |
|  | 200 | 412.2 | 1.19±0.05*** | - | - | 0.01±0.06 | - | 0.01±0.05 | - | - | - | - |
|  | 300 | 412.2 | 1.19±0.05*** | - | - | 0.01±0.05 | 0.01±0.05 | 0.01±0.05 | - | - | - | - |
|  | 400 | 412.2 | 1.19±0.05*** | - | - | 0.01±0.05 | 0.02±0.06 | 0.01±0.05 | - | - | - | - |
|  | 500 | 412.2 | 1.19±0.05*** | - | - | - | - | - | - | - | - | - |
