## Supplementary material for "Landscape composition mediates suppression of major pests by natural enemies in conventional cruciferous vegetables": Table S3

**Table S3. The list of the arthropod taxon and number of specimens collected in 2017 and 2018.**

| Order | Family | Species | 2017 | 2018 | Sum |
| --- | --- | --- | --- | --- | --- |
| Hemiptera | Aphididae | *Brevicoryne brassicae* (Linnaeus, 1758) | 26475 | 14236 | 40711 |
| Coleoptera | Chrysomelidae | *Phyllotreta vittula* (Redtenbacher, 1849) | 237 | 293 | 530 |
| Diptera | Agromyzidae | *Liriomyza sativae* Blanchard, 1938 | 913 | 917 | 1830 |
| Thysanoptera | Thripidae |  | 1705 | 1706 | 3411 |
| Lepidoptera | Plutellidae | *Plutella xylostella* (Linnaeus, 1758) | 3053 | 899 | 3952 |
| Hymenoptera | Aphelinidae |  | 164 | 163 | 327 |
| Hymenoptera | Bethylidae |  | 7 | 11 | 18 |
| Hymenoptera | Ceraphronidae |  | 47 | 190 | 237 |
| Hymenoptera | Chalcididae |  | 24 | 11 | 35 |
| Hymenoptera | Dryinidae |  | 2 | 0 | 2 |
| Hymenoptera | Eucharitidae |  | 5 | 2 | 7 |
| Hymenoptera | Diapriidaes |  | 101 | 375 | 476 |
| Hymenoptera | Encyrtidaes |  | 395 | 482 | 877 |
| Hymenoptera | Eulophidaes |  | 246 | 112 | 358 |
| Hymenoptera | Eupelmidae |  | 1 | 1 | 2 |
| Hymenoptera | Eurytomidae |  | 10 | 12 | 22 |
| Hymenoptera | Megaspilidae |  | 11 | 2 | 13 |
| Hymenoptera | Mymaridae |  | 33 | 37 | 70 |
| Hymenoptera | Perilampidae |  | 7 | 4 | 11 |
| Hymenoptera | Platygastridae |  | 183 | 116 | 299 |
| Hymenoptera | Proctotrupidae |  | 33 | 6 | 39 |
| Hymenoptera | Pteromalidae |  | 108 | 77 | 185 |
| Hymenoptera | Signiphoridae |  | 3 | 0 | 3 |
| Hymenoptera | Torymidae |  | 4 | 0 | 4 |
| Hymenoptera | Trichogrammatidae |  | 44 | 47 | 91 |
| Hymenoptera | Braconidae |  | 168 | 486 | 654 |
| Hymenoptera | Ichneumonidae |  | 209 | 129 | 338 |
| Coleoptera | Coccinellidae |  | 54 | 63 | 117 |
| Hymenoptera | Vespidae |  | 21 | 9 | 30 |
| Diptera | Syrphidae |  | 33 | 328 | 361 |
| Hymenoptera | Chrysididae |  | 3 | 4 | 7 |
| Neuroptera | Chrysopidae |  | 4 | 3 | 7 |
| Coleoptera | Staphylinidae |  | 906 | 3748 | 4654 |
| Dermaptera | Forficulidae |  | 79 | 35 | 114 |
| Araneae | Agelenidae | *Allagelena difficilis* (Fox, 1936) | 1 | 0 | 1 |
| Araneae | Araneidae | *Cyclosa japonica* Bösenberg & Strand, 1906 | 0 | 1 | 1 |
| Araneae | Araneidae | *Cyclosa sachikoae* Tanikawa, 1992 | 0 | 2 | 2 |
| Araneae | Araneidae | *Cyclosa argentata* Tanikawa & Ono, 1993 | 0 | 5 | 5 |
| Araneae | Araneidae | *Neoscona vigilans* (Blackwall, 1865) | 1 | 1 | 2 |
| Araneae | Atypidae | *Atypus* sp. | 2 | 0 | 2 |
| Araneae | Clubionidae | *Clubiona kulczynskii* Lessert, 1905 | 1 | 0 | 1 |
| Araneae | Gnaphosidae | *Gnaphosa microps* Holm, 1939 | 1 | 2 | 3 |
| Araneae | Gnaphosidae | *Odontodrassus hondoensis* (Saito, 1939) | 8 | 8 | 16 |
| Araneae | Gnaphosidae | *Zelotes sardus* (Canestrini, 1873) | 5 | 0 | 5 |
| Araneae | Gnaphosidae | *Zelotes tortuosus* Kamura, 1987 | 14 | 1 | 15 |
| Araneae | Hahniidae | *Hahnia* sp. | 5 | 10 | 15 |
| Araneae | Linyphiidae | *Caviphantes saxetorum* (Hull, 1916) | 0 | 1 | 1 |
| Araneae | Linyphiidae | *Erigone prominens* Bösenberg & Strand, 1906 | 439 | 368 | 807 |
| Araneae | Linyphiidae | *Gnathonarium dentatum* (Wider, 1834) | 61 | 143 | 204 |
| Araneae | Linyphiidae | *Walckenaeria* sp. | 0 | 2 | 2 |
| Araneae | Lycosidae | *Arctosa depectinata* (Bösenberg & Strand, 1906) | 2 | 1 | 3 |
| Araneae | Lycosidae | *Arctosa ipsa* (Karsch, 1879) | 0 | 0 | 0 |
| Araneae | Lycosidae | *Arctosa kwangreungensis* Paik & Tanaka, 1986 | 0 | 2 | 2 |
| Araneae | Lycosidae | *Pardosa distincta* (Blackwall, 1846) | 0 | 9 | 9 |
| Araneae | Lycosidae | *Pardosa laura* Karsch, 1879 | 19 | 18 | 37 |
| Araneae | Lycosidae | *Pirata subpiraticus* (Bösenberg & Strand, 1906) | 2 | 18 | 20 |
| Araneae | Lycosidae | *Trochosa aquatica* Tanaka, 1985 | 61 | 78 | 139 |
| Araneae | Lycosidae | *Wadicosa fidelis* (O.Pickard-Cambridge, 1872) | 2 | 19 | 21 |
| Araneae | Mysmenidae |  | 0 | 2 | 2 |
| Araneae | Nesticidae | *Nesticella mogera* (Yaginuma, 1972) | 15 | 19 | 34 |
| Araneae | Oxyopidae | *Hamataliwa* sp. | 2 | 1 | 3 |
| Araneae | Oxyopidae | *Oxyopes javanus* Thorell, 1887 | 0 | 1 | 1 |
| Araneae | Oxyopidae | *Oxyopes lineatipes* (C.L.Koch, 1847) | 3 | 1 | 4 |
| Araneae | Philodromidae | *Philodromus subaureolus* Bösenberg & Strand, 1906 | 0 | 1 | 1 |
| Araneae | Ctenidae | *Anahita* sp. | 1 | 0 | 1 |
| Araneae | Pisauridae | *Hygropoda higenaga* (Kishida, 1936) | 0 | 1 | 1 |
| Araneae | Psechridae | *Psechrus senoculatus* Yin, Wang & Zhang, 1985 | 0 | 1 | 1 |
| Araneae | Salticidae | *Myrmarachne japonica* (Karsch, 1879) | 0 | 1 | 1 |
| Araneae | Selenopidae | *Selenops* sp. | 2 | 0 | 2 |
| Araneae | Tetragnathidae | *Tetragnatha keyserlingi* Simon, 1890 | 4 | 13 | 17 |
| Araneae | Tetragnathidae | *Tetragnatha praedonia* L.Koch, 1878 | 1 | 0 | 1 |
| Araneae | Tetragnathidae | *Tetragnatha nitens* (Audouin, 1826) | 0 | 1 | 1 |
| Araneae | Theridiidae | *Cryptachaea blattea* (Urquhart, 1886) | 0 | 3 | 3 |
| Araneae | Theridiidae | *Emertonella taczanowskii* (Keyserling, 1886) | 0 | 2 | 2 |
| Araneae | Theridiidae | *Enoplognatha* sp. | 0 | 11 | 11 |
| Araneae | Theridiidae | *Meotipa pulcherrima* (Mello-Leitão, 1917) | 1 | 1 | 2 |
| Araneae | Theridiidae | *Platnickin* sp. | 0 | 2 | 2 |
| Araneae | Thomisidae | *Diaea subdola* O.Pickard-Cambridge, 1885 | 0 | 3 | 3 |
| Araneae | Thomisidae | *Thomisus* sp. | 0 | 1 | 1 |
| Araneae | Titanoecidae | *Nurscia albofasciata* (Strand, 1907) | 1 | 0 | 1 |
| Coleoptera | Carabidae | *Acupalpus elegans* (Dejean, 1829) | 2 | 0 | 2 |
| Coleoptera | Carabidae | *Acupalpus testaceus* (LeConte, 1844) | 2 | 9 | 11 |
| Coleoptera | Carabidae | *Agonum piceolum* (LeConte, 1879) | 12 | 64 | 76 |
| Coleoptera | Carabidae | *Amara littoralis* Dejean, 1828 | 1 | 4 | 5 |
| Coleoptera | Carabidae | *Anchomenus leucopus* Bates, 1873 | 6 | 11 | 17 |
| Coleoptera | Carabidae | *Anisodactylus punctatipennis* A.Morawitz, 1862 | 0 | 13 | 13 |
| Coleoptera | Carabidae | *Bembidion* sp. | 8 | 3 | 11 |
| Coleoptera | Carabidae | *Bradycellus* sp. | 2 | 1 | 3 |
| Coleoptera | Carabidae | *Chlaenius micans* (Fabricius, 1792) | 1 | 0 | 1 |
| Coleoptera | Carabidae | *Chlaenius posticalis* Motschulsky, 1854 | 0 | 9 | 9 |
| Coleoptera | Carabidae | *Colpodes* sp. | 0 | 2 | 2 |
| Coleoptera | Carabidae | *Colpodes* sp2. | 1 | 2 | 3 |
| Coleoptera | Carabidae | *Metacolpodes buchannani* (Hope, 1831) | 0 | 12 | 12 |
| Coleoptera | Carabidae | *Diplocheila zeelandica* (L.Redtenbacher, 1867) | 0 | 2 | 2 |
| Coleoptera | Carabidae | *Dolichus* sp. | 2 | 14 | 16 |
| Coleoptera | Carabidae | *Dolichus halensis* (Schaller, 1783) | 2 | 1 | 3 |
| Coleoptera | Carabidae | *Elaphropus* sp. | 3 | 5 | 8 |
| Coleoptera | Carabidae | *Harpalus anxius* (Duftschmid, 1812) | 0 | 3 | 3 |
| Coleoptera | Carabidae | *Harpalus calceatus* (Duftschmid, 1812) | 13 | 33 | 46 |
| Coleoptera | Carabidae | *Harpalus griseus* (Panzer, 1796) | 4 | 2 | 6 |
| Coleoptera | Carabidae | *Harpalus rufipes* DeGeer, 1774 | 4 | 5 | 9 |
| Coleoptera | Carabidae | *Harpalus servus* (Duftschmid, 1812) | 0 | 12 | 12 |
| Coleoptera | Carabidae | *Loxoncus cyanescens* (Hope, 1845) | 3 | 1 | 4 |
| Coleoptera | Carabidae | *Orthotrichus* sp. | 1 | 12 | 13 |
| Coleoptera | Carabidae | *Pentagonica daimiella* Bates, 1892 | 0 | 7 | 7 |
| Coleoptera | Carabidae | *Pheropsophus bimaculatus* (Linnaeus, 1771) | 0 | 1 | 1 |
| Coleoptera | Carabidae | *Pheropsophus jessoensis* A.Morawitz, 1862 | 4 | 9 | 13 |
| Coleoptera | Carabidae | *Platynus cincticollis* (Say, 1823) | 4 | 5 | 9 |
| Coleoptera | Carabidae | *Porotachys bisulcatus* (Nicolai, 1822) | 1 | 0 | 1 |
| Coleoptera | Carabidae | *Pterostichus anthracinus* (Illiger, 1798) | 0 | 15 | 15 |
| Coleoptera | Carabidae | *Pterostichus ovoideus* (Sturm, 1824) | 2 | 0 | 2 |
| Coleoptera | Carabidae | *Pterostichus strenuus* LeConte, 1853 | 5 | 12 | 17 |
| Coleoptera | Carabidae | *Pterostichus vernalis* (Panzer, 1796) | 7 | 0 | 7 |
| Coleoptera | Carabidae | *Stenolophus mixtus* (Herbst, 1784) | 41 | 5 | 46 |
| Coleoptera | Carabidae | *Stenolophus pseudoobockianus* Felix & Muilwijk, 2009 | 8 | 0 | 8 |
| Coleoptera | Carabidae | *Stenolophus quinquepustulatus* (Wiedemann, 1823) | 3 | 3 | 6 |
| Coleoptera | Carabidae | *Synuchus impunctatus* (Say, 1823) | 8 | 0 | 8 |
| Coleoptera | Carabidae | *Tachys gyotokuensis* Tanaka, 1956 | 26 | 10 | 36 |
| Coleoptera | Carabidae | *Trichotichnus longitarsis* A.Morawitz, 1863 | 22 | 76 | 98 |
| Coleoptera | Carabidae | *Trichotichnus vulpeculus* (Say, 1823) | 0 | 1 | 1 |
| Hymenoptera | Formcidae | *Pachycondyla javana* (Mayr, 1867) | 0 | 1 | 1 |
| Hymenoptera | Formcidae | *Brachyponera* sp. | 3 | 17 | 20 |
| Hymenoptera | Formcidae | *Pheidole noda* Smith, 1874 | 35 | 42 | 77 |
| Hymenoptera | Formcidae | *Nylanderia* sp. | 0 | 97 | 97 |
| Hymenoptera | Formcidae | *Monomorium chinense* Santschi, 1925 | 1 | 17 | 18 |
| Hymenoptera | Formcidae | *Pristomyrmex punctatus* (Smith, 1860) | 0 | 94 | 94 |
| Hymenoptera | Formcidae | *Polyrhachis dives* Smith, 1857 | 1 | 12 | 13 |
| Hymenoptera | Formcidae | *Tetramorium kraepelini* Forel, 1905 | 1 | 45 | 46 |
| Hymenoptera | Formcidae | *Aphaenogaster patruelis* Forel, 1886 | 8 | 0 | 8 |
| Hymenoptera | Formcidae | *Aphaenogaster patruelis* Forel, 1886 | 16 | 10 | 26 |
| Hymenoptera | Formcidae | *Crematogaster osakensis* Forel, 1900 | 0 | 22 | 22 |
| Hymenoptera | Formcidae | *Camponotus mitis* (Smith, 1858) | 0 | 4 | 4 |
| Hymenoptera | Formcidae | *Cardiocondyla venustula* Wheeler, 1908 | 0 | 5 | 5 |
| Hymenoptera | Formcidae | *Nylanderia vividula* (Nylander, 1846) | 0 | 12 | 12 |
| Hymenoptera | Formcidae | *Leptogenys* sp. | 0 | 4 | 4 |
| Hymenoptera | Formcidae | *Camponotus albosparsus* Bingham, 1903 | 0 | 1 | 1 |
| Hymenoptera | Formcidae | *Solenopsis invicta* Buren, 1972 | 0 | 102 | 102 |
| Hemiptera | Reduviidae | *Sirthenea flavipes* (Stål, 1855) | 2 | 0 | 2 |
| Hemiptera | Reduviidae | *Ectrychotes andreae* (Thunberg, 1784) | 0 | 2 | 2 |
| Hemiptera | Reduviidae | *Peirates turpis* Walker, 1873 | 0 | 1 | 1 |
| Hemiptera | Reduviidae | *Empicoris* sp. | 0 | 1 | 1 |
| Hemiptera | Reduviidae | *Peirates atromaculatus* (Stål, 1870) | 0 | 1 | 1 |
| Hemiptera | Reduviidae | *Sphedanolestes impressicollis* (Stål, 1861) | 2 | 0 | 2 |
| Hemiptera | Reduviidae | *Coranus* sp. | 1 | 0 | 1 |
